# Genome-wide association links candidate genes to fruit firmness, fruit flesh color, flowering time, and soluble solid content in apricot (*Prunus armeniaca*)

**DOI:** 10.1101/2020.04.16.044669

**Authors:** Filiz Ferik, Duygu Ates, Sezai Ercisli, Abdullah Erdogan, Emine Orhan, Muhammed Bahattin Tanyolac

## Abstract

Apricots originated from China, Central Asia and the Near East and arrived in Anatolia, and particularly in their second homeland of Malatya province in Turkey. Apricots are outstanding summer fruits, with their beautiful attractive colour, delicious sweet taste, aroma and high vitamin and mineral content. In the current study, a total of 259 apricot genotypes from different geographical origins in Turkey were used. Significant variations were detected in fruit firmness (FF), fruit flesh color (FFC), flowering time (FT), and soluble solid content (SSC). A total of 11,532 SNPs based on DArT were developed and used in the analyses of population structure and association mapping (AM). According to the STRUCTURE (v.2.2) analysis, the apricot genotypes were divided into three groups. The mixed linear model with Q and K matrixes were used to detect the associations between the SNPs and four traits. A total of 131 SNPs were associated with FF, FFC, and SSC. The results demonstrated that AM had high potential of revealing the markers associated with economically important traits in apricot.

## Introduction

Rosaceae is one of the most important fruit tree family from temperate regions, including apple, peach, strawberry, plum, almond, pear, European plum, and sweet cherry which are economically important fruit species (1)(2)(3). Apricot, *Prunus armeniaca* (Lam.), is also a member of this family and an important stone fruit with a total world production of about 3.8 million tons (4). The major apricot producing countries are Turkey (730.000 tons), Uzbekistan (662.123 tons), and Iran (306.115 tons) in the world (4).

Various parameters affect fruit quality in apricot (5)(6)(7). Consumer preferences are based on fruit quality which refers to sensorial properties, such as appearance, texture, taste and aroma, high nutritional components, chemical components, functional properties, and mechanical characteristics (8). The appearance of fruit is the main criterion for consumers, while a low level of sweetness and hard texture are undesired (9). Thus, fruit firmness and fruit flesh color are important for consumer satisfaction (10).

Another important parameter affecting fruit quality is soluble solid content (SSC), which includes sugars, organic acids, proteins, minerals, lipids, amino acids, and vitamins, and it is the main criteria that determines the taste, flavor and nutritional value of the fruit (11). In addition to being a delicious edible product, the fruit of apricot is also considered as functional due to its chemical ingredients (12)(13)(14)(15). The apricot fruit makes a significant contribution to human health through its phenolic compounds content with immune-stimulating, anti-inflammatory and antioxidants properties (16)(15).

One of the important traits for apricot producers is flowering time, which is a substantial agronomic trait with an impact on fruit and seed growth in temperate fruit tree species. In cold regions, early flowering individuals are damaged by late frost, while in warm regions, late flowering individuals could lead to some problems concerning leaf and flower bud break, resulting in a decrease in the amount of harvest (17). In addition, early flowering species have economic value in terms of early market prices.

Complex traits including most of the fruit quality properties are controlled by interacting genes called quantitative traits (18)(1). Therefore, the clarification of transmission of complex traits is one of the main topic for agricultural sciences (19). Quantitative trait locus (QTL) mapping is a prevalent method for the mapping of these kinds of traits and bases on biparental mapping populations (20). Constructing a new cross population is a tedious, time consuming and expensive process (21). Considering the long generation and juvenile period of fruit trees, it is more difficult to apply QTL mapping (19)(1). To date, a certain number of QTL maps have been constructed for apricot, such as flowering time (22)(23), resistance to sharka disease (24)(10)(25)(26), fruit quality traits (27)(28), chilling requirements (29), and tree architectural traits (28).

AM is an alternative method to pedigree-based QTL mapping (30) and uses natural populations, contrary to QTL mapping, to determine the correlations between phenotypes and genotypes (31). AM also utilizes historical recombination and natural variation as a basis and provides high map resolution in shorter time due to no requirement of developing a new cross population (30). This method has been previously employed for different fruit trees, such as peach (32)(33)(34)(35)(30), apple (36), almond (30), and apricot (29)(37).

The objective of this study was to investigate the associations between SNPs based on the diversity arrays technology (DArT) and the pomological traits of apricot, namely fruit firmness (FF), fruit flesh color (FFC), flowering time (FT), and SSC using 259 apricot genotypes.

## Materials and Methods

### Plant Material and DNA Isolation

A total of 259 apricot (*Prunus armeniaca* L.) genotypes, which were grown together at Malatya Apricot Research Institute in Turkey, were used in this study (S1Table). All genotypes had been planted (with 3 replications for each genotype) with eight-meter spacing between and within the rows in the experimental station of Malatya Apricot Research Institute, and all trees were 20 years old. Standard management practices concerning chemical fertilization, pruning, and disease control were being applied to the trees.

Young leaves were collected from each apricot genotype, cooled in liquid nitrogen, and stored at −80 °C for future analyses. The leaf samples were ground into small pieces with a tissue lyser (Technogen Co., Turkey). DNA extraction was carried out with 0.1 g samples of each individual following the protocol described by Deshmukh et al. (2007) with minor modifications. Tris-EDTA (TE) buffer (100 μl) was used to dissolve the extracted DNA. For the purification and quantification assessment of the isolated DNA, 1% agarose gel and a spectrophotometer (NanoDrop ND 1000) were used, respectively. After confirmation, the DNA samples were stored at −80 °C until they were used for SNP analyses.

### Pomological Evaluation

Forty fruits from each replication of genotype were randomly selected and harvested separately for each tree. FFC was measured with a Minolta Chroma Meter CR-400 (Minolta-Konica, Japan). For FT, observations were made by experts, and the first day of flowering was noted as FT. The juice of the 40 apricots was measured with a digital refractometer (Model RA-250HE Kyoto Electronics, Kyoto, Japan), and the SSC values were recorded in °Brix. FF was measured with an acoustic firmness sensor (Aweta BV, the Netherlands). These fruit traits were measured for two consecutive years (2016 and 2017).

### Variance Analysis

To define the variations in FF, FFC, FT and SSC among the 259 genotypes over two years (2016 and 2017), an analysis of variance (ANOVA) was performed using TOTEMSTAT software (39) according to the significance level of *P* ≤ 0.01. The variations were determined according to year (Y), genotype (G), and Y × G interactions.

### DArT analysis

The DArT analysis was performed as described by Nemli et al. (2017). The polymorphism information content (PIC) values represent the discrimination power of the markers. The PIC values were calculated for each marker according to Lynch and Ritland (1999) with the following equation: PIC=1-∑*p_i_^2^*, where *p_i_* demonstrates the proportion of the population with the *i*^th^ allele.

### Genetic Variation Analysis

STRUCTURE software (v.2.3.4), which is based on Bayesian modelling, was used to determine the population structure of 259 apricot genotypes (42). The software was run with a burn-in period of 10,000 and 10,000 Markov Chain Monte Carlo (MCMC) replications. Ten runs were performed for each number of populations (K), ranging from 1 to 10. The best number of subpopulations was determined with the Delta K (ΔK) value using STRUCTURE HARVESTER (43). For cross checking, the principle component analysis (PCA) was carried out with R Software [R statistical functions (R stats) and Gaussian mixture modelling for model-based clustering, classification, and density Estimation (mclust)], and a dendrogram tree was drawn with the same software with reference to Nei’s genetic distance (44).

### Association Mapping Analysis

TASSEL (v.5.2.3) software with a mixed linear model (MLM (K+Q) model) was used for the detection of the associations between DNA markers and pomological traits (FF, FFC, FT, and SSC) (45). The relative kinship matrix, which shows the genetic relationships between the individuals, was calculated by TASSEL (v.5.0) based on the centered IBS method (45). The Q matrix was obtained from STRUCTERE software at the ΔK = 3 value. The associations between the SNP markers and pomological traits were visualized as Manhattan plots in R software with the “qqman” package. The designation of significant markers was performed in the same software with the false discovery rate (FDR) (46) and Bonferroni correction (47) being calculated separately for each pomological trait (FF, FFC, FT and SSC). Furthermore, the quantile-quantile (Q-Q) plots were visualized with the same software.

### Identification of candidate genes

The sequences of the SNP markers associated with FF, FFC, and SSC were analyzed to determine the functions of the candidate genes using the Phtozome (v12.1) database.

## Results

### Phenotypic Variation

In the present study, FF, FFC, FT, and SSC were measured for two years (2016-2017), and showed normal distribution (S1-4 Fig). These findings indicated the importance of the genetic background of each genotype for the *Prunus* phenotyping profile. The minimum, maximum and mean values of each year showed high consistency, and no significant differences were observed between the years (2016 and 2017) for the mean values. The minimum, maximum and mean values of all phenotypic traits are presented in S2 Table.

The mean values of four traits (FF, FFC, FT and SSC) only slightly differed between the two years (2016 and 2017). However, there were fourfold differences between the SSC and FFC values obtained from 2016 and 2017 (Table 1). FT ranged from 95 to 125 days with a mean value of 114.2 days in 2016, and it ranged from 95 to 126 days with a mean value of 112 days in 2017 (Table 1). FF varied between 0.1 N and 9.60 N in 2016 and 0.03 N and 6.62 N in 2017 (Table 1). There was a nearly 90-fold difference in the ranges obtained for FF from the two years. The individuals showing the highest and lowest average values are listed in S2 Table.

**Table 1.** Minimum, maximum and mean values of all phenotypic traits

| Trait | 2016 |  |  | 2017 |  |  |
| --- | --- | --- | --- | --- | --- | --- |
|  | Min | Max | Mean | Min | Max | Mean |
| FF | 0.1 N | 9.60 N | 1.86 N | 0.03 N | 6.62 N | 1.71 N |
| FFC | 11.6° | 46.72° | 34.04° | 13.99° | 46.77° | 34.12° |
| FT | 95 days | 125 days | 114.24 days | 95 days | 126 days | 112 days |
| SSC | 8.22°Brix | 29.92°Brix | 15.28°Brix | 8.23°Brix | 32.37°Brix | 16.27°Brix |

The results of the correlation analysis showed no significant correlation between the four traits (FF, FFC, FT and SSC) (S3 Table). The results of ANOVA are presented in S4 Table. ANOVA demonstrated significant variations according to year, genotype, and year × genotype interactions for all apricot traits at the P ≤ 0.01 significance level (S4 Table).

### Population Structure Analysis

A total of 24,864 SNP markers were generated from the DArT analysis, and after filtering the missing data (max 5% missing data, Marker Allele Frequency (MAF)>0.5), 11,532 high-quality SNP markers were obtained. The PIC value was 0.77, ranging between 0.05 and 0.99. These markers were assigned to the related scaffolds and were used in the STRUCTURE (v.2.2) analysis. This analysis was performed for K from 1 to 10, and the peak was observed at K = 3 according to the ΔK computation data. The STRUCTURE results showed that the 259 genotypes were divided into three main populations: namely POPI (red), POPII (green) and POPIII (blue) (S5 Fig). All these genotypes were also further divided into three groups according to Nei’s genetic distance analysis: The first group consisted of Geno 185 (Nigde – Turkey) and Geno 186 (Malatya-Turkey), the second comprised Geno 38 (Siverek/Urfa – Turkey), Geno 230 (USA) and Geno 255 (Russia), and the third contained the remaining 254 genotypes. These results indicate that the genotypes used in this study were not clustered according to their geographical origin. In addition, PCA revealed three groups for the population used in this study (S6 Fig), confirming the results obtained from STRUCTURE (S7 Fig) and Nei’s genetic distance analyses (S4 Fig).

The expected heterozygosity and fixation index (*Fst*) are parameters that explain the heterozygosity level of a population. In this study, the expected heterozygosity was determined as 0.20 for Cluster 1, 0.06 for Cluster 2, and 0.11 for Cluster 3, with a mean value of 0.12. On the other hand, the *Fst* value varied between 0.14 and 0.81 with a mean value of 0.55, representing a high genetic variation level for the population.

### AM Analysis

AM analyses were carried out for four pomological traits (FF, FFC, FT and SSC) using TASSEL (v.5.2.3) software and the MLM (Q+K) model in two consecutive years (2016 and 2017). These analyses detected a large number of associations related to the pomological traits. FDR and Bonferroni corrections were applied to eliminate the false positives among the associations. Eventually, 131 SNP markers were found to be associated with three traits (FF, FFC, and SSC). Among these associations, three, 57 and 71 SNPs were associated with FFC, FF and SSC respectively.

A total of 88 and 228 SNPs were associated with FF in 2016 and 2017, respectively (FDR correction applied, −log_10_*P* ≥ 2.90 for 2016 and ≥ 2.78 for 2017), and 57 of these markers were common for both 2016 and 2017 (S5 and S6 Tables and Fig 1). Most of the significant SNPs for FF were detected in 2017 but not in 2016 (S5 and S6 Tables and Fig 1).

**Fig 1.**
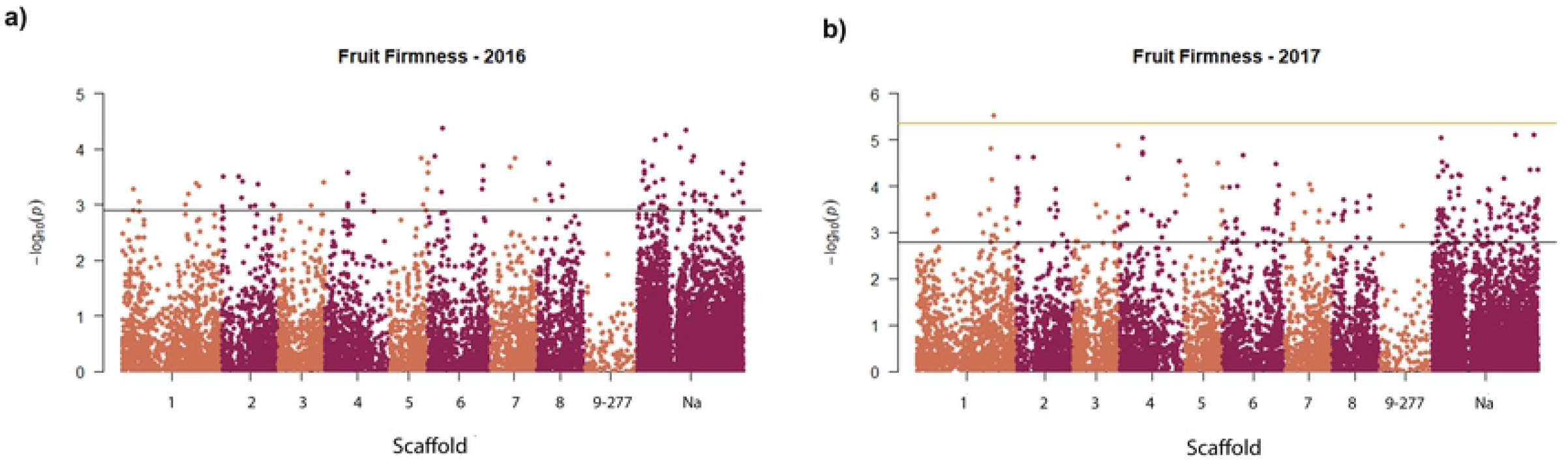
Manhattan plots of fruit firmness for the two years

For FFC, three SNPs (−log_10_ *P* value is ≥ 3.27, FDR correction applied) and 13 SNPs (−log_10_ *P* ≥ 3.10, FDR correction applied) were associated with FFC in 2016 and 2017, respectively, and three of these SNPs (SNP 4257, SNP 17194 and SNP 22875) was common for both years (S5 Table and Fig 2).

**Fig 2.**
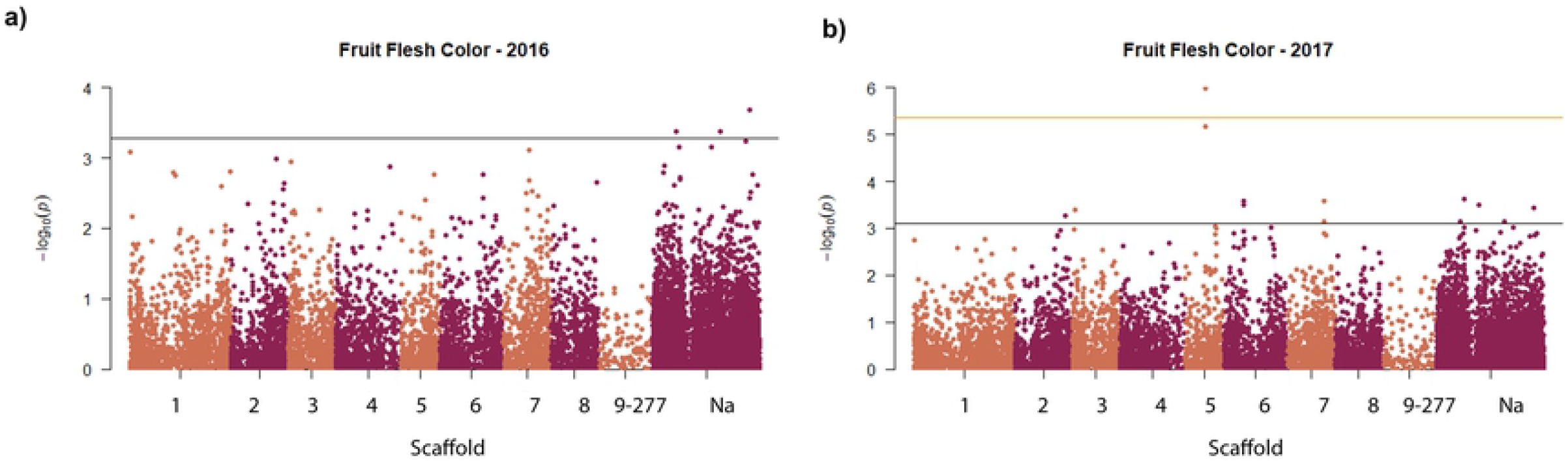
Manhattan plots of fruit flesh color for the two years

The marker-trait association analysis for FT revealed that it was associated with 10 SNPs (FDR correction applied, −log_10_ *P* ≥3.28) in 2016 and 22 SNPs (FDR correction applied, −log_10_ *P* ≥3.06) in 2017. However, none of these SNPs was commonly seen in both years (S5 Table and Fig 3).

**Fig 3.**
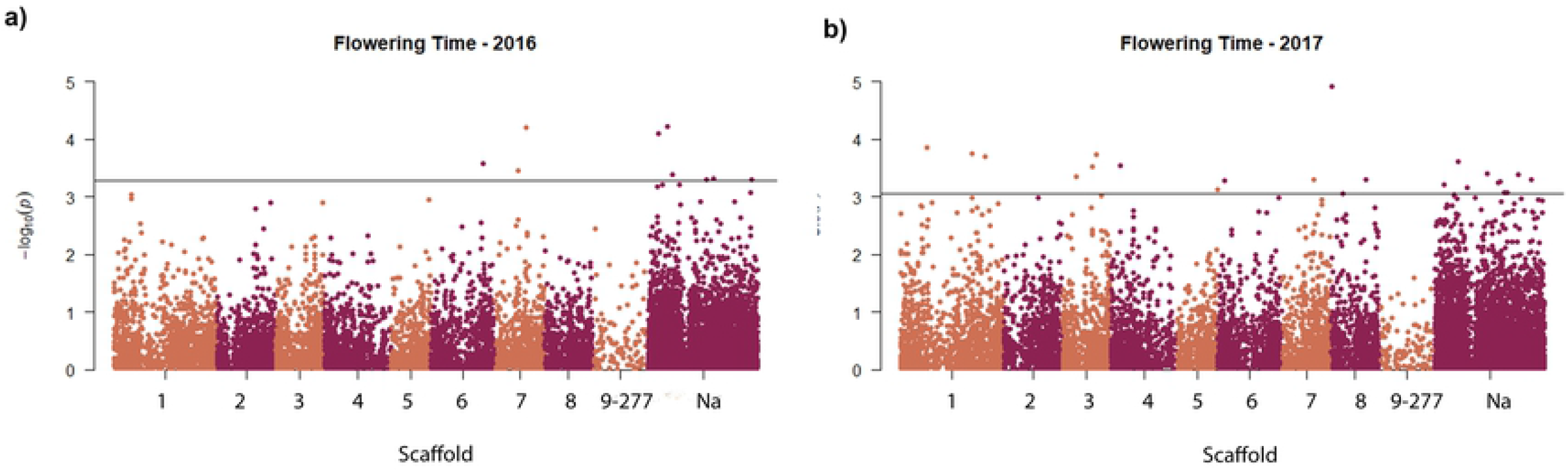
Manhattan plots of flowering time for the two years

For SSC, 167 SNPs (−log_10_ *P ≥* 2.82, FDR correction applied) and 352 SNPs (−log_10_ *P ≥* 2.72, FDR correction applied) were found related in 2016 and 2017, respectively. Of these SNPs, 71 were detected in both years (Fig 4 and S5 and S7 Tables). The *P* values presenting the significance level of the associations between the markers and pomological traits are given in Q-Q plots in S8 Fig.

**Fig 4.**
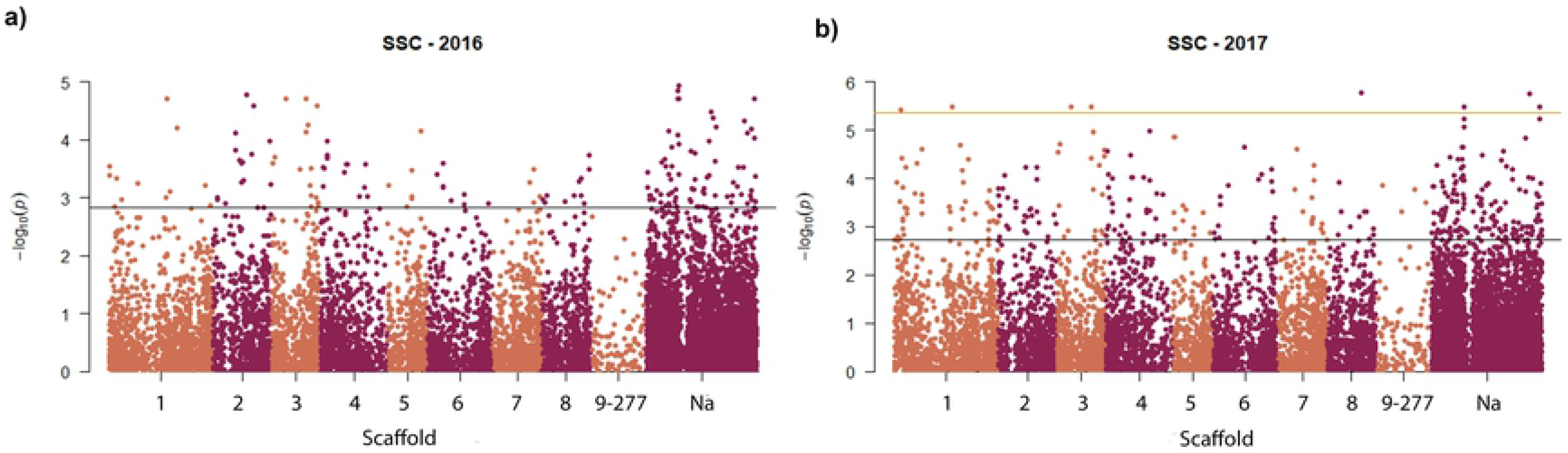
Manhattan plots of solid soluble content for the two years

### Identification of Candidate Genes

A total number of 30 putative candidate genes were found to be related to the SNPs associated with FF and SSC (S8 Table). SNPs which were associated with the FFC trait did not show similarity to any of the putative candidate genes. For the SSC of the apricots, the following proteins and enzymes related to putative candidate genes showed homology with SNPs (given in parentheses): putative 3,4-dihydroxy-2-butanone kinase (SNP526), putative leucine-rich repeat receptor-like protein kinase (SNP1023), pentatricopeptide repeat-containing protein (SNP1482), probable LRR receptor-like serine/threonine-protein kinase (SNP1494), vignain-like (SNP2823), transcription termination factor MTERF6 (SNP3309), LRR receptor-like serine/threonine-protein kinase GSO2 (SNP3842), WAT1-related protein At5g64700 (SNP4443 and SNP20435), cell division cycle 20.2, cofactor of APC complex-like (SNP4948), long-chain acyl-CoA synthetase8 (SNP5158), calcium-transporting ATPase 12 plasma membrane-type-like (SNP5444), probable LRR receptor-like serine/threonine-protein kinase At4g36180 (SNP13398), peroxidase 5 (SNP14227), UDP-glycosyltransferase TURAN (SNP15803), tRNA (guanine(37)-N1)-methyltransferase 1 (SNP16184), transcription factor TFIIIB component B’’ homolog (SNP16422), receptor like protein 30-like (SNP17431), putative disease resistance protein (SNP19126), putative pre-16S rRNA nuclease (SNP19568), DNA-directed RNA polymerase III subunit (SNP 20753), DEAD-box ATP-dependent RNA helicase 3 (SNP22678), putative pentatricopeptide repeat-containing protein (SNP23251), S locus F-box protein f (SLFf) gene, partial cds; and Sf-RNase and S haplotype-specific F-box protein f (SFBf) genes, complete cds (SNP23325), putative disease resistance RPP13-like protein 1 (SNP23732), TMV resistance protein N-like (SNP23833), putative disease resistance protein (SNP23953), LRR receptor-like serine/threonine-protein kinase GSO2 (SNP24483), and receptor-like protein 12 (SNP24611) (S8 Table).

## Discussion

### Phenotypic Variation

Fruit quality parameters are of prime importance for both consumers and growers. Among these parameters, FT is the most widely studied physical attribute in apricot due to its economic importance related to early market prices (28)). In the present study, FT was detected between 95 and 126 days (Table 1). Among the genotypes studied, Geno115 was the earliest cultivar with 95 days. In previous studies, FT was reported to range from 59 to 84 days (23) and 55 to 78 days (28). The variation that was detected in the present study for FT was greater compared to the literature and can be attributed to the different genetic origins of the cultivars.

SSC is one of the main criteria affecting fruit taste, and SSC value greater than 12°Brix indicates good gustative quality (10). In the present study, SSC was measured between 8.22 and 32.36 °Brix, most genotypes (243 genotypes) had a value over 12°Brix, there was nearly a fourfold difference in the range (Table 1). In different studies, SSC was detected as 8.73 to 17.80 °Brix (27), 10.6 to 16.2 °Brix (10) and 6.2 to 19.5 °Brix (28). In our study, a larger variation was found in SSC compared to previous studies. This wide range of phenotypic discrepancy indicates the genotypic variation level of the apricot genotypes used in the study.

FFC and FF are two most important sensorial properties that affect consumer preferences at the purchase step. In the present study, FFC was measured between 11.6° and 46.77°, and there was nearly a fourfold difference in the range (Table1). In previous studies, FFC was measured from 67° to 100° (27) and 70° to 94.7° (10). In the current study, the range of FF was obtained as 0.03 to 9.60 N. In the literature, FF was reported to vary between 24.9 and 62.2 N (10) and 15 and 50 N (27). Both FFC and FF ranges measured in the current study were quite different from those of previous studies.

### Population Structure

In genetic mapping studies, associations between DNA markers and traits are affected by the type and number of markers (48). The use of a high number of markers leads to high genome coverage, and therefore high-throughput systems gain importance. In the current study, DArT, a high-throughput system, was used and a total of 20,264 SNP markers were developed. These markers were used in population diversity. This high number of markers provided high genome coverage.

The STRUCTURE analysis was used for the identification of the population structure of the 259 apricots investigated in the current study. These genotypes were divided into three main populations (S2 Fig). PCA (S3 Fig) and dendrogram (S4 Fig) analyses confirmed the number of populations. These genotypes were not divided into populations according to their geographical origin. For example, the genotypes that originated from Turkey, Spain, Italy, Poland, Armenia, France, USA, and Hungary were included in the same (third) group (S4 Fig). The reason for this result could be the complex breeding history of these genotypes. In particular, the use of cultivars with different histories in introgression and intercrossing processes may have led to this situation (48). In addition, humans move plants from one geographic realm to another, which results in confusion concerning the origin of the plants (49). In a previous AM study on apricot, 72 genotypes were used, and the genotypes were divided into two main groups (37). The reason for the lower number of subpopulations in the current study could be the use of a population with a narrow genetic basis. Although the previous authors also selected the genotypes from different countries, they may have used those of the same origin.

In the present study, the mean *Fst* value was detected as 0.55 and the mean expected heterozygosity was 0.12, indicating the presence of a high genetic variation in the population structure. These findings also support the idea that DArT systems develop a large number of SNP markers distributed along the apricot genome. In previous studies, the *Fst* value was reported to be 0.51 (50) and 0.16 (51), and the expected heterozygosity value was 0.82 (50) and 0.29 (51). These differences between previous studies in terms of diversity may be due to the number and type of markers utilized, and genotyping being undertaken in distinct locations (52).

### Association Mapping Analysis

AM is a powerful technique which is based on the accumulation of genetic variability through evolution in natural populations to identify DNA markers based on the association between genetic markers and phenotypes (53). AM analyses are used as a rapid and efficient alternative to linkage mapping analyses (30) for detecting the associations between traits and markers; therefore, these analyses are widely employed in the mapping of economically important traits in many crop species (54)(55). In the present study, K and Q matrixes were used to correct the population structure in the MLM (Q+K) model included in AM analyses. This model effectively eliminates possible false positives with random and fixed effects according to Henderson’s notation (56). In addition, FDR and Bonferroni corrections were applied to eliminate spurious associations (56). To date, no AM study has been undertaken to reveal the associations between SNP markers and pomological traits (FF, FFC, FT and SSC). However, an association map was constructed by Mariette et al. (2016) to identify the SNP markers that controlled resistance to plum pox virus in apricot. Furthermore, Olukolu (2010) constructed an AM on the chilling requirements of apricot. Apart from these studies, there are only a limited number of association studies on the economically important traits of the other members of the *Rosaceae* family (57)(33)(58)(30)(59)(55).

In the present study, a total of 131 SNPs were found significantly associated with three pomological traits (FF, FFC and SSC) of the apricot genotypes via AM analyses (S5-7 Tables and Fig 1–4). Three SNPs (SNP 4257, SNP17194 and SNP 22875) were found associated with FFC and 57 SNPs were found associated with FF (S5-7 Tables and Fig1–4). A total of 71 SNPs were associated with SSC in two consecutive years (2016 and 2017) (S5 and S7 Tables). In previous studies, (27) and (28) used control cross populations of apricot and found one QTL each that was related to SSC. The number of significant markers identified in the present study was higher than previously reported (27)(28) which may be related to the type of population investigated. In mapping studies, natural populations provide higher genome coverage and mapping resolution with regard to wide genotypical variations (56). Another reason for our high number of SNPs may be the use of DArT to produce the markers. This technology is known to produce a high number of SNP markers, and thus provide high genome coverage.

### Identification of Candidate Genes

In the present study, 30 putative candidate genes showed homology with the sequences of SNPs associated with FF and SSC (S8 Table). Among these, transcription termination factor MTERF6 plays an important role in plastid development on *Arabidopsis thaliana* (60). WAT1-related protein is located on cell wall and responsible for transmembrane transporter activity (61). Long-chain acyl-CoA synthetase 8 is very important for lipid metabolism (62). UDP-glycosyltransferase TURAN is one of the responsible enzyme for development of pollen tube. LRR receptor-like serine/threonine-protein kinase (GSO2) and probable LRR receptor-like serine/threonine-protein kinase (GSO1) together play a role in root growth and the growth of epidermal surface in embryos and cotyledons in *Arabidopsis* (63). Putative pentatricopeptide repeat-containing protein is a member of the pentatricopeptide repeat (PPR) protein family and is involved in the organellar RNA metabolism (64). Calcium-transporting ATPase 12, plasma membrane-type-like has a function in calcium-transporting ATPase activity and calmodulin binding (65). tRNA (guanine(37)-N1)-methyltransferase 1 is responsible for the methylation of cytoplasmic and mitochondrial tRNAs in the N1 position of guanosine-37 *Arabidopsis thaliana* (66). TMV resistance protein N-like provides resistance to tobacco mosaic virus in plants (67). Putative 3,4-dihydroxy-2-butanone kinase plays a role in ATP binding (68). Pentatricopeptide repeat-containing protein is a required protein for the intergenic processing between chloroplast rsp7 and ndhB transcripts (69).

## Conclusions

This is the first AM study that presents the associations between SNPs based on the DArT technology and economically important traits (FF, FFC, FT and SSC) in apricot. Large variations were determined for these four traits. The results of this study highlight the importance of using populations with wide variations in AM studies. AM revealed significant associations for FF, FFC and SSC. The SNPs identified in the study can be used in future breeding programs for marker-assisted selection in apricot. On the other hand, the genotypes with the shortest FT can be used as parents in developing earlier cultivars combined with other desirable traits.

## Author Contribution Statement

FF: Association map analysis, writing manuscript; DA: writing manuscript, data obtaining; SE: PI of the projects; AE: obtained phenotyping data, EO: statistical analysis, data obtaining; MBT: corresponding author, conceived and designed research.

## Acknowledgements

The study was financially supported by the Scientific and Technological Research Council of Turkey (TUBITAK) with the project number TUBITAK-215O445.

## Conflict of interest

The authors declare no competing interests.

## Supporting Information

**S1 Fig** Histogram showing the distribution of fruit firmness in 2016 and 2017

**S2 Fig** Histogram showing the distribution of fruit flesh color in 2016 and 2017

**S3 Fig** Histogram showing the distribution of flowering time in 2016 and 2017

**S4 Fig** Histogram showing the distribution of solid soluble content in 2016 and 2017

**S5 Fig** STRUCTURE plot of 259 individuals by 20,264 SNPs at Δ*K* = 3

**S6 Fig** PCA analysis of 259 individuals

**S7 Fig** Dendrogram tree analysis based on Nei’s genetic distance

**S8 Fig** Q-Q plots of fruit firmness, fruit flesh color, flowering time, and solid soluble content

**S1 Table** List of 259 apricot genotypes

**S2 Table** Individuals with the highest and lowest average values of investigated traits

**S3 Table** Correlation analysis results

**S4 Table** Summary of ANOVA for fruit firmness, fruit flesh color, flowering time, and solid soluble content

**S5 Table** Number of SNP markers associated with the investigated traits according to the FDR correction threshold

**S6 Table** List of SNP markers identified as significantly associated with fruit firmness

**S7 Table** List of SNP markers identified as significantly associated with SSC

**S8 Table** Details of identification of putative candidate gene associations

## References

1. Meneses C, Orellana A. Using genomics to improve fruit quality. Vol. 46, Biological Research. Sociedad de Biología de Chile; 2013. p. 347–52.

2. Agric For TJ, Butiuc-keul A, Coste A, Farkas A, Cristea V, Isac V, et al. Turkish Journal of Agriculture and Forestry Molecular characterization of apple (Malus × domestica Borkh.) genotypes originating from three complementary conservation strategies. journals.tubitak.gov.tr [Internet]. [cited 2020 Apr 12]; Available from: http://journals.tubitak.gov.tr/agriculture/

3. Agric For TJ, Soysal D, Demirsoy L, Macit İ, Lang G, Demirsoy H. Turkish Journal of Agriculture and Forestry The applicability of new training systems for sweet cherry in Turkey. journals.tubitak.gov.tr [Internet]. [cited 2020 Apr 12]; Available from: https://www.mgm.gov.tr/2015

4. FAOSTAT [Internet]. [cited 2020 Apr 13]. Available from: http://www.fao.org/faostat/en/#data/QC

5. Bassi D, Science RS-A in H, 1990 undefined. Evaluation of fruit quality in peach and apricot. JSTOR [Internet]. [cited 2020 Apr 12]; Available from: https://www.jstor.org/stable/42883062?casa_token=0H2zwEno-VYAAAAA:SP5eu8deSgubahtbgu-G-Ob2F9gDlL4cTLojrtZuQeEbMpuibQJPMPbUO99-wh6-ZuPRMsR-ON36WQzmMEmgG-1WZ6rA97CPgZagmDw_4msQhIn-2yo

6. Altindag M, Sahin M, Esitken A, Ercisli S, Control MG-B, 2006 undefined. Biological control of brown rot (Moniliana laxa Ehr.) on apricot (Prunus armeniaca L. cv. Hacıhaliloğlu) by Bacillus, Burkholdria, and Pseudomonas application under. Elsevier [Internet]. [cited 2020 Apr 12]; Available from: https://www.sciencedirect.com/science/article/pii/S1049964406001204

7. Halász J, Pedryc A, Ercisli S, … KY-J of the, 2010 undefined. S-genotyping supports the genetic relationships between Turkish and Hungarian apricot germplasm. journals.ashs.org [Internet]. [cited 2020 Apr 12]; Available from: https://journals.ashs.org/jashs/view/journals/jashs/135/5/article-p410.xml

8. Abbott JA. Quality measurement of fruits and vegetables [Internet]. Vol. 15, Postharvest Biology and Technology. 1999 [cited 2020 Apr 12]. Available from: https://www.sciencedirect.com/science/article/pii/S0925521498000866

9. Defilippi BG, Predieri S, Infante R. Apricot (Prunus armeniaca L.) quality and breeding perspectives SEE PROFILE [Internet]. 2014 [cited 2020 Apr 12]. Available from: www.world-food.net

10. Soriano JM, Vera-Ruiz EM, Vilanova S, Martínez-Calvo J, Llácer G, Badenes ML, et al. Identification and mapping of a locus conferring plum pox virus resistance in two apricot-improved linkage maps. Tree Genet Genomes. 2008 Jul;4(3):391–402.

11. Akin E, Karabulut I, Chemistry AT-F, 2008 undefined. Some compositional properties of main Malatya apricot (Prunus armeniaca L.) varieties. Elsevier [Internet]. [cited 2020 Apr 12]; Available from: https://www.sciencedirect.com/science/article/pii/S0308814607008473

12. Dragovic-Uzelac V, Levaj B, Mrkic V, chemistry DB-F, 2007 undefined. The content of polyphenols and carotenoids in three apricot cultivars depending on stage of maturity and geographical region. Elsevier [Internet]. [cited 2020 Apr 12]; Available from: https://www.sciencedirect.com/science/article/pii/S0308814606002962

13. Drogoudi PD, Vemmos S, Pantelidis G, Petri E, Tzoutzoukou C, Karayiannis I. Physical Characters and Antioxidant, Sugar, and Mineral Nutrient Contents in Fruit from 29 Apricot (Prunus armeniaca L.) Cultivars and Hybrids. J Agric Food Chem [Internet]. 2008 Nov 26 [cited 2020 Apr 12];56(22):10754–60. Available from: http://pubs.acs.org

14. Leccese A, Bureau S, Reich M, Renard MGCC, Audergon JM, Mennone C, et al. Pomological and nutraceutical properties in apricot fruit: Cultivation systems and cold storage fruit management. Plant Foods Hum Nutr. 2010;65(2):112–20.

15. Hegedus A, Engel R, Abrankó L, Balogh E, Blázovics A, Hermán R, et al. Antioxidant and Antiradical Capacities in Apricot (Prunus armeniaca L.) Fruits: Variations from Genotypes, Years, and Analytical Methods. J Food Sci. 2010 Nov;75(9).

16. Madrau MA, Piscopo A, Sanguinetti AM, Del Caro A, Poiana M, Romeo F V., et al. Effect of drying temperature on polyphenolic content and antioxidant activity of apricots. Eur Food Res Technol. 2009 Jan;228(3):441–8.

17. Fan S, Bielenberg DG, Zhebentyayeva TN, Reighard GL, Okie WR, Holland D, et al. Mapping quantitative trait loci associated with chilling requirement, heat requirement and bloom date in peach (Prunus persica). New Phytol [Internet]. 2010 Mar 1 [cited 2020 Apr 12];185(4):917–30. Available from: http://doi.wiley.com/10.1111/j.1469-8137.2009.03119.x

18. Illa E, Eduardo I, Audergon JM, Barale F, Dirlewanger E, Li X, et al. Saturating the Prunus (stone fruits) genome with candidate genes for fruit quality. Mol Breed. 2011 Dec;28(4):667–82.

19. Neale DB, Savolainen O. Association genetics of complex traits in conifers. Elsevier [Internet]. [cited 2020 Apr 12]; Available from: www.sciencedirect.com

20. Würschum T. Mapping QTL for agronomic traits in breeding populations. Vol. 125, Theoretical and Applied Genetics. 2012. p. 201–10.

21. Shulaev V, Korban SS, Sosinski B, Abbott AG, Aldwinckle HS, Folta KM, et al. Multiple models for Rosaceae genomics. Vol. 147, Plant Physiology. American Society of Plant Biologists; 2008. p. 985–1003.

22. Campoy JA, Ruiz D, Egea J, Rees DJG, Celton JM, Martínez-Gómez P. Inheritance of Flowering Time in Apricot (Prunus armeniaca L.) and Analysis of Linked Quantitative Trait Loci (QTLs) using Simple Sequence Repeat (SSR) Markers. Plant Mol Biol Report. 2011 Jun;29(2):404–10.

23. Dirlewanger E, Quero-Garcia J, Heredity LLD-, 2012 undefined. Comparison of the genetic determinism of two key phenological traits, flowering and maturity dates, in three Prunus species: peach, apricot and sweet cherry. nature.com [Internet]. [cited 2020 Apr 12]; Available from: https://www.nature.com/articles/hdy201238

24. Lambert P, Dicenta F, Rubio M, Audergon JM. QTL analysis of resistance to sharka disease in the apricot (Prunus armeniaca L.) “Polonais” × “Stark Early Orange” F1 progeny. Tree Genet Genomes. 2007 Oct;3(4):299–309.

25. Marandel G, Salava J, Abbott A, Candresse T, Decroocq V. Quantitative trait loci meta-analysis of Plum pox virus resistance in apricot (Prunus armeniaca L.): New insights on the organization and the identification of genomic resistance factors. Mol Plant Pathol. 2009 May;10(3):347–60.

26. Dondini L, Lain O, Vendramin V, Rizzo M, Vivoli D, Adami M, et al. Identification of QTL for resistance to plum pox virus strains M and D in Lito and Harcot apricot cultivars. Mol Breed. 2011 Jan;27(3):289–99.

27. Salazar JA, Ruiz D, Egea J, Martínez-Gómez P. Transmission of Fruit Quality Traits in Apricot (Prunus armeniaca L.) and Analysis of Linked Quantitative Trait Loci (QTLs) Using Simple Sequence Repeat (SSR) Markers. Plant Mol Biol Report. 2013 Dec;31(6):1506–17.

28. Socquet-Juglard D, Christen D, Devènes G, Gessler C, Duffy B, Patocchi A. Mapping Architectural, Phenological, and Fruit Quality QTLs in Apricot. Plant Mol Biol Report. 2013;31(2):387–97.

29. Olukolu BA, Trainin T, Fan S, Kole C, Bielenberg DG, Reighard GL, et al. Genetic linkage mapping for molecular dissection of chilling requirement and budbreak in apricot (*Prunus armeniaca* L.). Gulick P, editor. Genome [Internet]. 2009 Oct [cited 2020 Apr 12];52(10):819–28. Available from: http://www.nrcresearchpress.com/doi/10.1139/G09-050

30. Font i Forcada C, Velasco L, Socias i Company R, Fernández i Martí Á. Association mapping for kernel phytosterol content in almond. Front Plant Sci. 2015 Jul 9;6(JULY).

31. Kaya HB, Cetin O, Kaya HS, Sahin M, Sefer F, Tanyolac B. Association Mapping in Turkish Olive Cultivars Revealed Significant Markers Related to Some Important Agronomic Traits. Biochem Genet. 2016 Aug 1;54(4):506–33.

32. Aranzana MJ, Abbassi EK, Howad W, Arús P. Genetic variation, population structure and linkage disequilibrium in peach commercial varieties. BMC Genet. 2010 Jul 20;11.

33. Cao K, Wang L, Zhu G, Fang W, Chen C, Luo J. Genetic diversity, linkage disequilibrium, and association mapping analyses of peach (Prunus persica) landraces in China. Tree Genet Genomes. 2012;8(5):975–90.

34. Font i Forcada C, Oraguzie N, Igartua E, Moreno MÁ, Gogorcena Y. Population structure and marker-trait associations for pomological traits in peach and nectarine cultivars. Tree Genet Genomes. 2013;9(2):331–49.

35. Picañol R, Eduardo I, Aranzana MJ, Howad W, Batlle I, Iglesias I, et al. Combining linkage and association mapping to search for markers linked to the flat fruit character in peach. Euphytica. 2013;190(2):279–88.

36. Cevik V, Ryder CD, Popovich A, Manning K, King GJ, Seymour GB. A FRUITFULL-like gene is associated with genetic variation for fruit flesh firmness in apple (Malus domestica Borkh.). Tree Genet Genomes. 2010 Jan;6(2):271–9.

37. Mariette S, Wong Jun Tai F, Roch G, Barre A, Chague A, Decroocq S, et al. Genome-wide association links candidate genes to resistance to Plum Pox Virus in apricot (Prunus armeniaca). New Phytol. 2016 Jan 1;209(2):773–84.

38. Deshmukh V, Thakare P, … UC-EJ of, 2007 undefined. A simple method for isolation of genomic DNA from fresh and dry leaves of Terminalia arjuna (Roxb.) Wight and Arnot. scielo.conicyt.cl [Internet]. [cited 2020 Apr 12]; Available from: https://scielo.conicyt.cl/scielo.php?pid=S0717-34582007000300014&script=sci_arttext&tlng=n

39. Acikgoz N, Ilker E, University AG-E, TOTEM undefined, Izmir undefined, 2004 undefined. Assessment of biological research on the computer.

40. Agric For TJ, Nemli S, Kaygisiz Aşçioğul T, Ateş D, Eşiyok D, Bahattin TANYOLAÇ M. Turkish Journal of Agriculture and Forestry Diversity and genetic analysis through DArTseq in common bean (Phaseolus vulgaris L.) germplasm from Turkey. [cited 2020 Apr 13]; Available from: http://journals.tubitak.gov.tr/agriculture/

41. Lynch M, Ritland K. Estimation of Pairwise Relatedness With Molecular Markers [Internet]. Genetics Soc America. 1999 [cited 2020 Apr 13]. Available from: https://www.genetics.org/content/152/4/1753.short

42. Pritchard J, Stephens M, Genetics PD-, 2000 undefined. Inference of population structure using multilocus genotype data. Genet Soc Am [Internet]. [cited 2020 Apr 12]; Available from: https://www.genetics.org/content/155/2/945.full-text.pdf+html

43. Evanno G, Regnaut S, Goudet J. Detecting the number of clusters of individuals using the software STRUCTURE: A simulation study. Mol Ecol. 2005 Jul;14(8):2611–20.

44. Nei M. Genetic Distance between Populations. Am Nat. 1972 May;106(949):283–92.

45. Bradbury P, Zhang Z, Kroon D, … TC-, 2007 undefined. TASSEL: software for association mapping of complex traits in diverse samples. academic.oup.com [Internet]. [cited 2020 Apr 13]; Available from: https://academic.oup.com/bioinformatics/article-abstract/23/19/2633/185151

46. Benjamini Y, Hochberg Y. Controlling the False Discovery Rate: A Practical and Powerful Approach to Multiple Testing. J R Stat Soc Ser B. 1995 Jan;57(1):289–300.

47. carboni CB-S in onore del professore salvatore ortu, 1935 undefined. Il calcolo delle assicurazioni su gruppi di teste. ci.nii.ac.jp [Internet]. [cited 2020 Apr 12]; Available from: https://ci.nii.ac.jp/naid/20001029336/

48. Mather D, Hayes P, Chalmers K, Eglinton J, Matus I, Richardson K, et al. Use of SSR Marker Data to Study Linkage Disequilibrium and Population Structure in Hordeum vulgare: Prospects for Association Mapping in Barley. Czech Journal of Genetics and Plant Breeding; 2004.

49. Pyšek P, Richardson DM, Rejmánek M, Webster GL, Williamson M, Kirschner J. Alien plants in checklists and floras: towards better communication between taxonomists and ecologists. Taxon [Internet]. 2004 Feb [cited 2019 Dec 24];53(1):131–43. Available from: http://doi.wiley.com/10.2307/4135498

50. Khan M, Maghuly F, … EB-F-S, 2008 undefined. Genetic diversity and population structure of apricot (Prunus armeniaca L.) from Northern Pakistan using Simple Sequence Repeats. degruyter.com [Internet]. [cited 2020 Apr 13]; Available from: https://www.degruyter.com/downloadpdf/j/sg.2008.57.issue-1-6/sg-2008-0024/sg-2008-0024.xml

51. Tian-Ming H, Xue-Sen C, Zheng X, Jiang-Sheng G, Pei-Jun L, Wen L, et al. Using SSR markers to determine the population genetic structure of wild apricot (Prunus armeniaca L.) in the Ily Valley of West China. Genet Resour Crop Evol. 2007 May 26;54(3):563–72.

52. Ates D, Asciogul TK, Nemli S, Erdogmus S, Esiyok D, Tanyolac MB. Association mapping of days to flowering in common bean (Phaseolus vulgaris L.) revealed by DArT markers. Mol Breed. 2018 Sep 1;38(9):1–14.

53. Ozkuru E, Ates D, Nemli S, Erdogmus S, Karaca N, Yilmaz H, et al. Genome-wide association studies of molybdenum and selenium concentrations in C. arietinum and C. reticulatum seeds. Mol Breed. 2019 Mar 1;39(3).

54. Khazaei H, Podder R, Caron C, … SK-T plant, 2017 undefined. Marker–trait association analysis of iron and zinc concentration in lentil (Lens culinaris Medik.) seeds. dl.sciencesocieties.org [Internet]. [cited 2020 Apr 12]; Available from: https://dl.sciencesocieties.org/publications/tpg/abstracts/10/2/plantgenome2017.02.0007

55. Font I. Forcada C, Guajardo V, Chin-Wo SR, Moreno MÁ. Association mapping analysis for fruit quality traits in prunus persica using SNP markers. Front Plant Sci. 2019 Jan 17;9.

56. Cappa EP, El-Kassaby YA, Garcia MN, Acuña C, Borralho NMG, Grattapaglia D, et al. Impacts of population structure and analytical models in genome-wide association studies of complex traits in forest trees: A case study in Eucalyptus globulus. PLoS One. 2013 Nov 25;8(11).

57. Ganopoulos I V., Kazantzis K, Chatzicharisis I, Karayiannis I, Tsaftaris AS. Genetic diversity, structure and fruit trait associations in Greek sweet cherry cultivars using microsatellite based (SSR/ISSR) and morpho-physiological markers. Euphytica. 2011 Sep;181(2):237–51.

58. Dhanapal AP, Crisosto CH. Association genetics of chilling injury susceptibility in peach (Prunus persica (L.) Batsch) across multiple years. 3 Biotech. 2013 Dec;3(6):481–90.

59. Cao K, Zhou Z, Wang Q, Guo J, Zhao P, … GZ-N, et al. Genome-wide association study of 12 agronomic traits in peach. nature.com [Internet]. [cited 2020 Apr 12]; Available from: https://www.nature.com/articles/ncomms13246

60. Romani I, Manavski N, Morosetti A, Tadini L, Maier S, Kühn K, et al. A Member of the Arabidopsis Mitochondrial Transcription Termination Factor Family Is Required for Maturation of Chloroplast Transfer RNA Ile (GAU). Am Soc Plant Biol [Internet]. 2010 [cited 2020 Apr 12]; Available from: www.plantphysiol.org/cgi/doi/10.1104/pp.15.00964

61. Kovi M, Amdahl H, Alsheikh M, reports OR-S, 2017 undefined. De novo and reference transcriptome assembly of transcripts expressed during flowering provide insight into seed setting in tetraploid red clover. nature.com [Internet]. [cited 2020 Apr 12]; Available from: https://www.nature.com/articles/srep44383

62. Zhao L, Haslam T, Sonntag A, … IM-P and C, 2019 undefined. Functional Overlap of Long-Chain Acyl-CoA Synthetases in Arabidopsis. academic.oup.com [Internet]. [cited 2020 Apr 12]; Available from: https://academic.oup.com/pcp/article-abstract/60/5/1041/5304662

63. Racolta A, Bryan AC, Tax FE. The receptor-like kinases GSO1 and GSO2 together regulate root growth in arabidopsis through control of cell division and cell fate specification. Dev Dyn. 2014 Feb;243(2):257–78.

64. Williams PM, Barkan A. A chloroplast-localized PPR protein required for plastid ribosome accumulation. Plant J. 2003 Dec;36(5):675–86.

65. Song P, Chen X, Wu B, Gao L, Zhi H, Cui X. Identification for soybean host factors interacting with P3N-PIPO protein of Soybean mosaic virus. Acta Physiol Plant. 2016 Jun 1;38(6).

66. Lee C, Kramer G, Graham DE, Appling DR. Yeast mitochondrial initiator tRNA is methylated at guanosine 37 by the Trm5-encoded tRNA (guanine-N1-)-methyltransferase. J Biol Chem. 2007 Sep 21;282(38):27744–53.

67. Hehl R, Faurie E, Whitham S, Baker B, Hesselbach J, Salamini F, et al. TMV resistance gene N homologues are linked to Synchytrium endobioticum resistance in potato. Theor Appl Genet. 1999;98(3–4):379–86.

68. Herz S, Kis K, Bacher A, Phytochemistry FR-, 2002 undefined. A tomato enzyme catalyzing the phosphorylation of 3, 4-dihydroxy-2-butanone. Elsevier [Internet]. [cited 2020 Apr 12]; Available from: https://www.sciencedirect.com/science/article/pii/S0031942202000560

69. Barkan A, Small I. Pentatricopeptide Repeat Proteins in Plants. Annu Rev Plant Biol. 2014 Apr 29;65(1):415–42.

